# Detection of Gray Mold Infection in Plants Using a Multispectral Imaging System

**DOI:** 10.1101/2020.04.23.051300

**Authors:** Clifton G. Scarboro, Stephanie M. Ruzsa, Colleen J. Doherty, Michael W. Kudenov

**Affiliations:** Optical Sensing Lab, North Carolina State University, Dept. of Elect. and Comp. Engineering, 2410 Campus Shore Drive, Raleigh, NC 27606; North Carolina State University, Dept. of Molecular and Structural Biochemistry, 120 W Broughton Drive, Raleigh, NC 27607

**Keywords:** multispectral imaging, hyperspectral imaging, reflectance indexing, gray mold, plant pathology, disease detection

## Abstract

Gray mold disease caused by the fungus *Botrytis cinerea* damages many crop hosts worldwide and is responsible for heavy economic losses. Early diagnosis and detection of the disease would allow for more effective crop management practices to prevent outbreaks in field or greenhouse settings. Furthermore, having a simple, non-invasive way to quantify the extent of gray mold disease is important for plant pathologists interested in quantifying infection rates. In this paper, we design and build a multispectral imaging system for discriminating between leaf regions, infected with gray mold, and those that remain unharmed on a lettuce (*Lactuca spp.)* host. First, we describe a method to select two optimal (high contrast) spectral bands from continuous hyperspectral imagery (450-800 nm). We then built a system based on these two spectral bands, located at 540 and 670 nm. The resultant system uses two cameras, with a narrow band-pass spectral filter mounted on each, to measure the multispectral reflectance of a lettuce leaf. The two resulting images are combined using a normalized difference calculation that produces a single image with high contrast between the leaves’ infected and healthy regions. A classifier was then created based on the thresholding of single pixel values. We demonstrate that this simple classification produces a true positive rate of 95.25% with a false positive rate of 9.316%.

## 1. Introduction

*Botrytis cinerea* is an airborne plant pathogen that causes losses by means of gray mold disease in over 200 different species of crops worldwide, with the most damage being seen in dicotyledonous hosts [1]. Typical symptoms of the disease observed in plant leaves and fruits include soft rots followed by the appearance of gray masses of conidia [1]. A color image of various Salinas and US96 lettuce (*Lactuca sativa*) leaves with severely advanced infection is shown in Figure 1.

**Figure 1.**
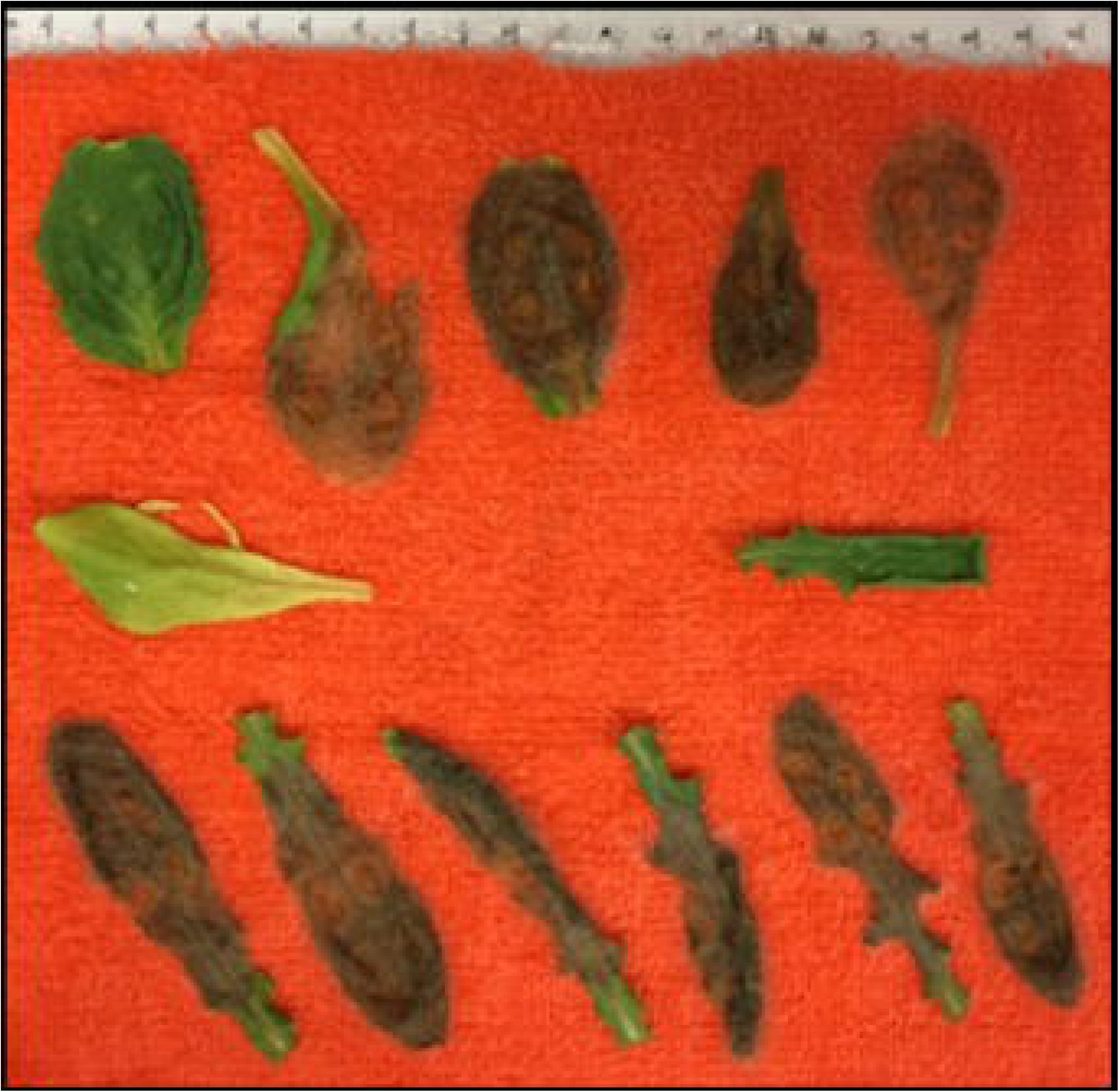
Salinas (top row) and US96 (bottom row) lettuce leaves seven days after inoculation with *Botrytis cinerea* in a controlled temperature chamber. The middle row of leaves contains uninoculated control Salinas (left) and US96 (right) leaves.

Gray mold is also to be a major threat to greenhouse-grown plants, where control of the disease is usually challenging and expensive [2]. Careful application of fungicides would allow for effective management of gray mold when growing plants. Using fewer well-timed sprays of fungicide is more efficient than more frequent applications through harvest due to the possibility of promoting resistant populations of *B. cinerea* [3]. An imaging system that can highlight areas of gray mold infection in plant leaves with accuracy and consistency would allow for quick and easy assessment of the disease on a large number of plants at once so that growers can make informed decisions on spraying fungicide. In addition, as demonstrated here, such an imaging system provides a useful tool for plant pathologists studying the spread of gray mold. The ability to consistently and quantitatively detect the first signs of *B. cinerea* and to monitor disease progression in a leaf can provide detailed trait information to researchers comparing treatment methods or investigating genotype-driven differences in susceptibility.

Several studies have addressed gray mold diagnosis and detection in plants. Traditionally, fungal disease-causing pathogens have been detected by visual examination of symptoms on the plant [4]. This involves interpreting disease symptoms such as blight, rotting, spots, and wilting using available guidelines and standards for assessment [4]. Some drawbacks to this method are that its effectiveness is subject to an examiners judgement and therefore produces variable results. Scoring by eye is also time-consuming and can be difficult to scale for more high-throughput analysis or high-resolution temporal analysis throughout the infection progression. Molecular quantification based on quantitative polymerase chain reaction of quantifying fungal conidiospores offer increased throughput [5]. However, these methods are destructive and therefore, the same individual plant cannot be monitored across time. Additional detection methods include plating and culturing plant pathogens for identification using microscopy techniques [6], but again these approaches are challenging to scale for high-throughput, high-resolution temporal analysis. Less time-consuming methods of detecting the extent of infection in a plant population involves the use of spectroscopic and imaging techniques. Fungal diseases can alter leaf morphology, and therefore their light reflectance, which can be distinguished using hyperspectral or multispectral imaging [7]. Fahrentrapp *et. al.* demonstrated that measuring the reflectance of a leaf at narrow-band near-infrared (NIR) wavelengths was sufficient to distinguish between a leaf’s healthy and infected areas by using a linear regression model [7]. An issue with the model used is that it requires taking into account the relative position between the leaf and detector, as well as the lighting conditions. Wu et al. [8] used a hyperspectral visible near-infrared (VNIR) spectroradiometer to measure reflectance intensities of healthy and *B. cinerea*-inoculated eggplant leaves. By performing a principal component analysis on the collected data and training a back-propagation neural network on this data, the authors were able to predict infection with 85% accuracy before infection symptoms were visible. The results relied upon collecting data in a controlled environment where ambient lighting was limited. However, while such VNIR approaches may offer improved early detection, they are more expensive and complex to deploy in automated imaging applications. Two-band detection methods have also been tested. Mo *et al.* found optimal wavebands at 552 nm and 701 nm in a two-band ratio for detecting discoloration in Salinas lettuce leaves with over 99% accuracy; however, its application towards *Botrytis* detection was not studied [9].

In this paper, we demonstrate a process for selecting optimal spectral bands to be integrated in a two-band multispectral camera imaging system and evaluate its performance, for the purposes of detecting *B. cinerea* infection on *L. sativa* leaves to monitor disease progression. In section 2, we outline the theory needed to assess the hyperspectral data, while in section 3, we describe the experimental setup used to acquire our hyperspectral and multispectral data. Section 4 describes the results of our timelapse experiments, while section 5 ends with a conclusion.

## 2. Materials & Methods

### 2.1 Spectral Band Optimization

Hyperspectral and multispectral imaging devices have been widely used for measuring irradiance spectra of a scene with varying spectral resolution, range, and acquisition methods. They measure light collected from a scene as a function of two spatial dimensions and one spectral dimension. The main distinction between a hyperspectral and multispectral system is the quantity of spectral bands measured by the device. There are a variety of methods for acquiring this three-dimensional data that are mostly described by the way they discriminate the spatial and spectral dimensions [10]. Examples of spatial data acquisition methods include whiskbroom (point-scanning) and pushbroom (line-scanning) systems, as well as “snapshot” two-dimensional imagers [10]. Some ways these devices acquire spectral data include spectral filtering, dispersive elements like prisms or diffraction gratings, and interferometric Fourier transform methods [10].

To discover the optimal spectral bands, we use a prism-based pushbroom scanning hyperspectral imaging camera that has been calibrated per Ref. [11] to provide comprehensive lettuce irradiance spectra. Relative reflectance was calculated to quantify both healthy and infected lettuce leaf tissues, such that

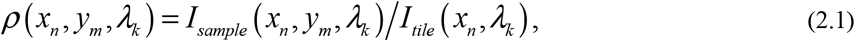

where *I*_*sample*_ is the intensity measured from the lettuce, *I*_*tile*_ is the downwelling intensity measured from a white (99% reflectivity) spectralon tile illuminated by the source, *x*_*n*_ and *y*_*m*_ are the *x* and *y* coordinates corresponding to the discrete integer spatial position *n* and *m*, respectively, and *λ*_*k*_ is the *k*^th^ wavelength element in the datacube. For visualization and labeling purposes, 2-dimensional (2D) panchromatic imagery was generated by integrating our 3-dimensional (3D) data cube along the spectral axis for each image slice, such that

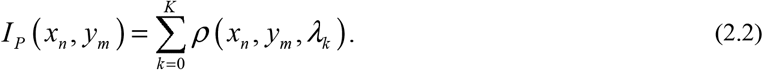

After reflectivity has been calculated, we leveraged graphical reflectance indexing to establish a simple analysis of the scene’s spectral measurements. By taking radiance measurements at specific spectral components, correlations between features of the scene can be made. A common example of this is the normalized difference vegetation index (NDVI) which takes the difference of reflectance of two spectral bands in the near infrared (NIR) and red regions and divides by their sum [12]. Generally, the contrast, *ν*, from such two-band calculations for arbitrary wavelengths, *λ*_1_ and *λ*_2_, are described by

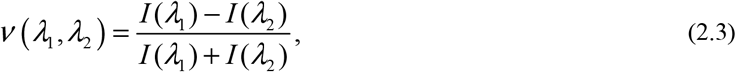

where the intensity measurements *I* are implicitly dependent upon *x*_*n*_ and *y*_*m*_ for clarity. This procedure minimizes the effect of lighting and atmospheric conditions on measurements to enable more accurate comparison [13]. Also, image data from two spectral bands can be condensed into one single image that is produced from the contributions of both bands.

The goal for a system implementing reflectance indices is to select two spectral bands where the contrast is optimized between two characteristics of interest, *a* and *b* (*e.g.*, symptomatic versus healthy areas of lettuce leaves). Using data from the hyperspectral reflectance plots, a heat map, which is defined by

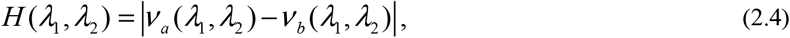

was obtained by sweeping through each combination of two spectral bands, *λ*_1_ and *λ*_2_ as inputs to Eq. (2.3) to calculate the contrasts, *ν*_*a*_ and *ν*_*b*_, associated with the characteristics, *a* and *b*. By locating local maxima in *H*, the optimal spectral bands can be determined for use in a multispectral camera system.

### 2.2 Experimental Optimization

To calculate *v* over many combinations of spectra, a pushbroom hyperspectral imaging camera was used to collect a three-dimensional spatial and spectral data cube over a visible to near-infrared spectral range with wavelength resolution spanning 0.5 to 2.4 nm for 500 to 800 nm light. Lettuce leaves of the variety Black Seed Simpson were inoculated with *B. cinerea* which spread throughout the leaves over the course of a few days. The leaves were placed on a damp paper towel, inside a 10 in x 10 in bioassay dish (Corning). Treatment leaves were inoculated with *B. cinerea* suspended in Potato dextrose broth (PDB). Control leaves, in the same tray, were mock-inoculated with PDB alone. When the disease symptoms were significant (about 50 percent coverage of the leaf area) on most of the inoculated lettuce leaves, the tray was imaged by the pushbroom camera, and a tungsten light source was used to illuminate the sample leaves. The pushbroom camera and tungsten light source setup is presented in Figure 2. A tungsten halogen lamp was configured to illuminate the target at a normal angle of incidence from the leaf’s surface normal. Our pushbroom spectral camera was positioned 40 cm away, yielding a spatial resolution of approximately 6.5 mm. Light entered the objective lens, slit, and collimator where it was dispersed using a prism. Dispersed light was then imaged onto a focal plane array (FPA) using a reimaging lens and raw data were calibrated in accordance to Ref. [11]. The FPA measures an image with a spatial dimension along the slit and a spectral dimension caused by the dispersive prism. In order to acquire the other spatial dimension required for a two-dimensional image, the entire imaging system is mounted on a rotating platform that moves while the camera acquires successive frames that form the three-dimensional data cube, *I*_*sample*_.

**Figure 2.**
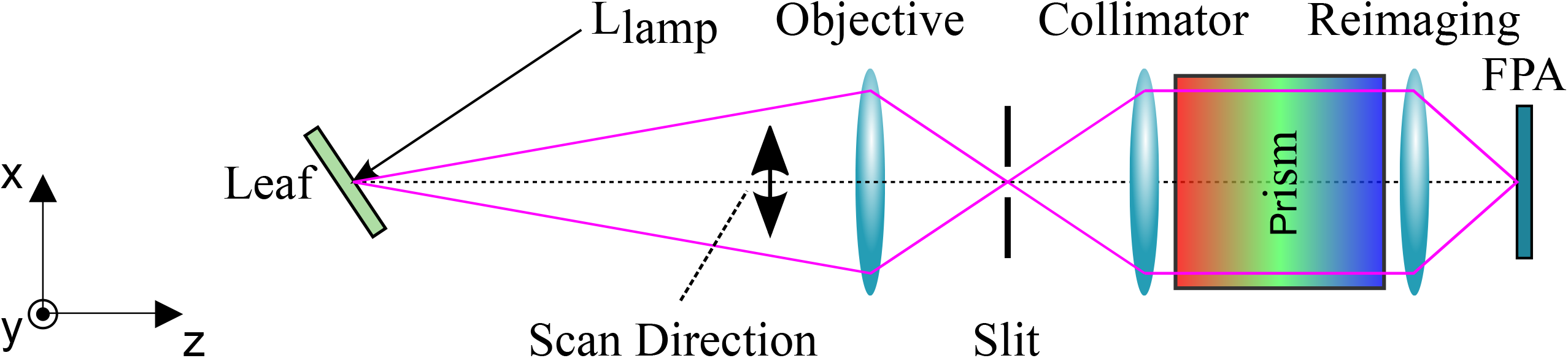
Hyperspectral pushbroom camera for preliminary characterization of lettuce infected with *Botrytis cinerea.*

After acquiring the hyperspectral datacubes, two-dimensional images of the lettuce were obtained using Eq. (2.2). Infected and control (healthy) tissues, from inoculated and control leaves, respectively, were manually labeled by defining regions of interest (ROIs). Each ROI was then spatially averaged to create a normalized reference spectrum, which are depicted in Figure 3. Generally, the normalized reflectivity is the same except around 540 nm and 720 nm. Our reflectance measurements presented in Figure 3 are supported by reflectivity measurements taken by Simko *et al.* for lettuce tissue in fresh and decayed conditions [14].

**Figure 3.**
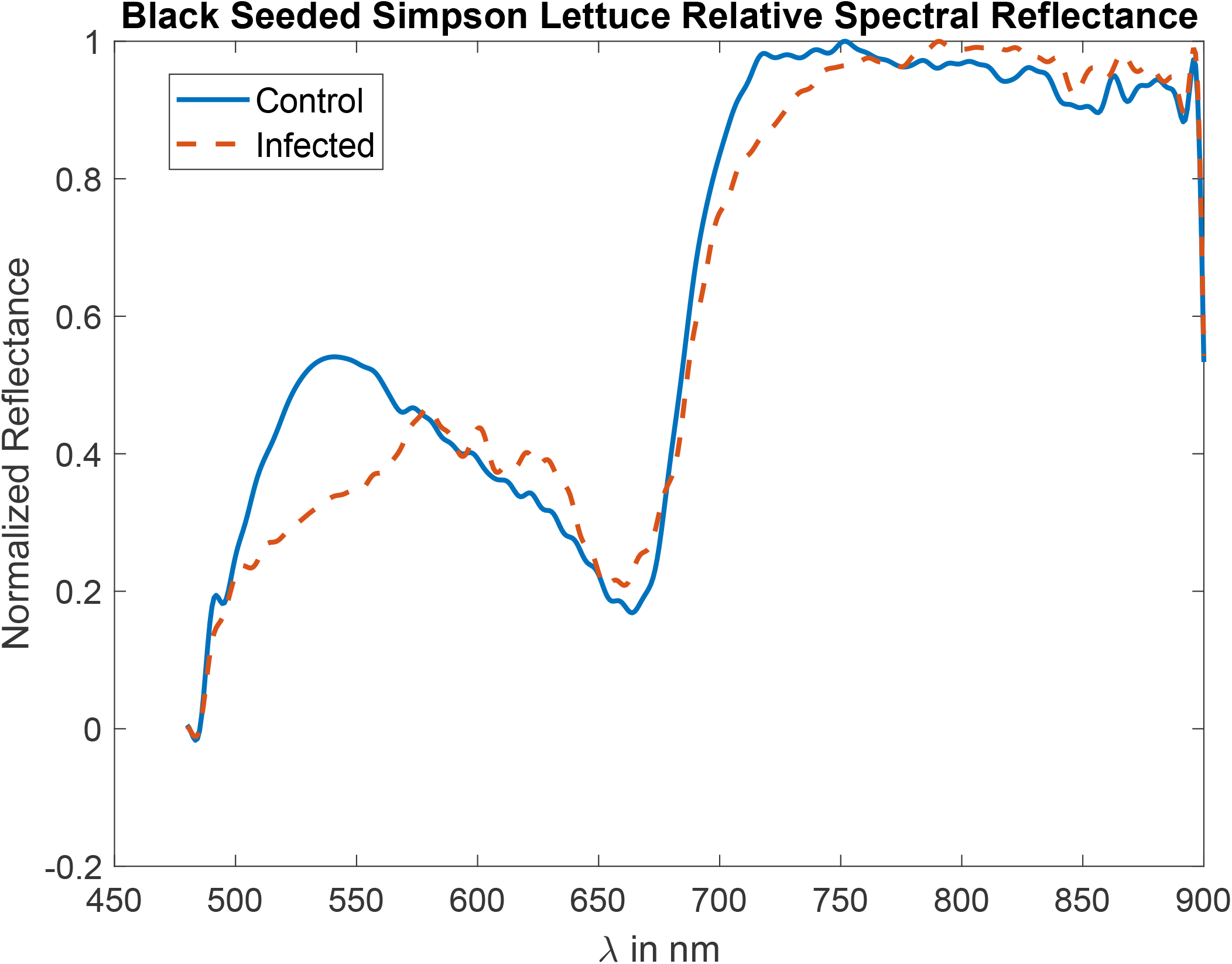
Normalized spectral response measured for infected and control lettuce leaves of the Black Seeded Simpson variety.

The reference spectra were further used to produce the heat map in Figure 4. It should be noted that the heat map is symmetric about the diagonal due to the same spectral bands used in the calculation from Eq. (2.4). From these results, the highest contrast is located at *λ*_1_ = 670 nm and *λ*_2_ = 530 nm.

**Figure 4.**
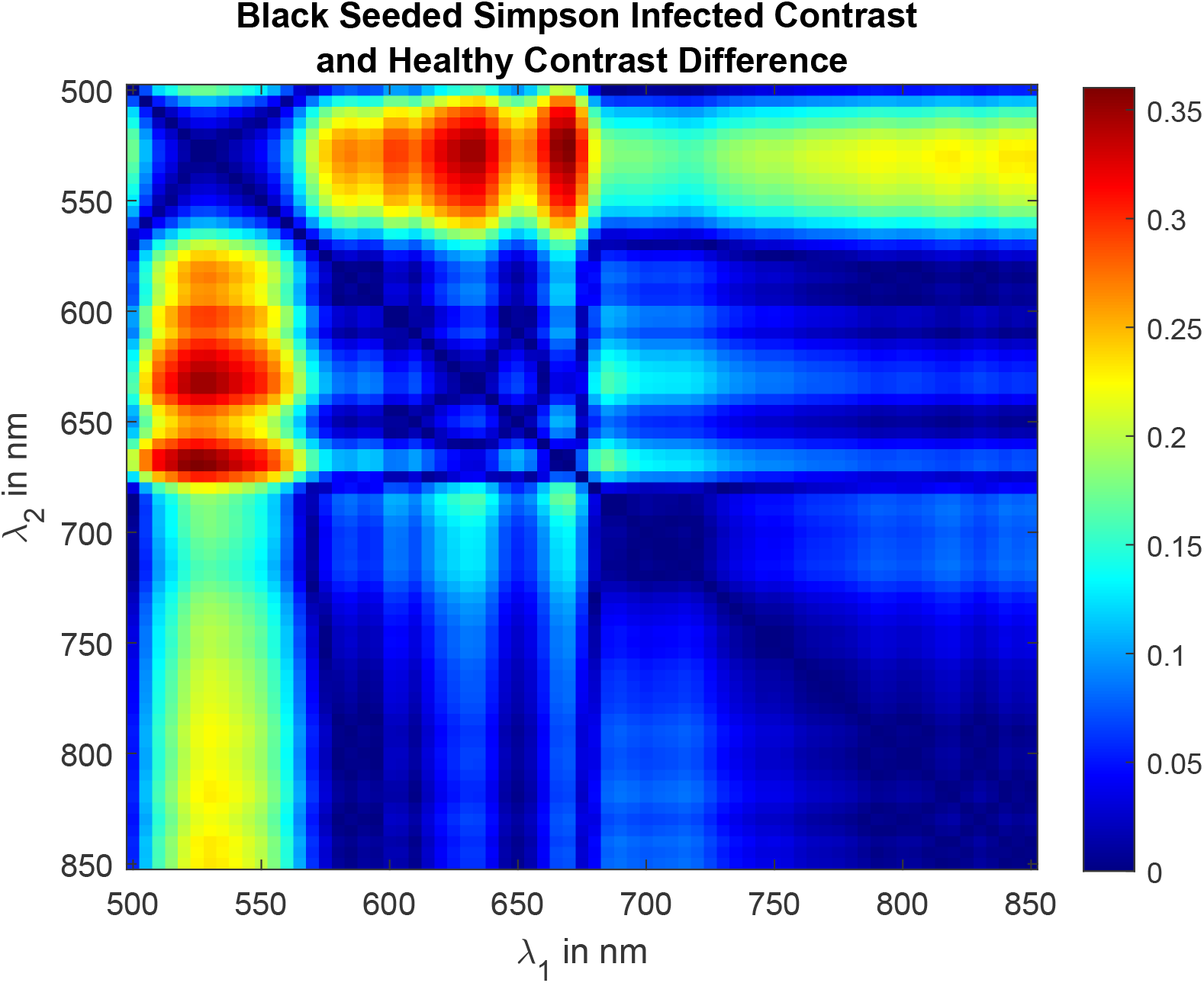
The absolute difference in contrasts of two spectral bands measured on healthy and infected lettuce areas for the Black Seeded Simpson type.

Using this information, we selected two spectral band-pass filters for use in a dual-camera multispectral system. The filters selected had center pass-bands at 540 nm and 670 nm filter and a 10 nm full width at half maximum spectral bandwidth. Note that one filter (540 nm) deviates slightly from our calculated maximum due to off-the-shelf filter availability at the time of purchase; however, this is only expected to reduce *v* by 2.8%.

A schematic of the 2-band imaging system is depicted in Figure 5 (a). Light from the sample first enters a beam splitter (BS), which directs light from a scene to each camera. Transmitted or reflected light from the BS then enters the spectral band-pass filters BP1 or BP2, respectively, before forming an image by L1 or L2 onto FPA1 or FPA2, respectively. The FPA consisted of two BlackFly 1.3 megapixel, 30 frames per second USB3 monochrome cameras. Both cameras were configured to take images of the scene simultaneously. A photograph of the completed experimental setup is depicted in Figure 5 (b). Since this system was used to take lettuce measurements over several days, a white LED source was used that does not radiate as much heat as the tungsten lamp in order to lessen environmental stress on the lettuce leaves. In addition, experiments were performed inside of a clear plexiglass container prevent the spread of *B. cinerea* to the surrounding environment. Since the imaging system was located outside of the plexiglass barrier, glare off its surface was reduced by placing the LED off-axis in an illumination geometry similar to darkfield microscopy [15].

**Figure 5.**
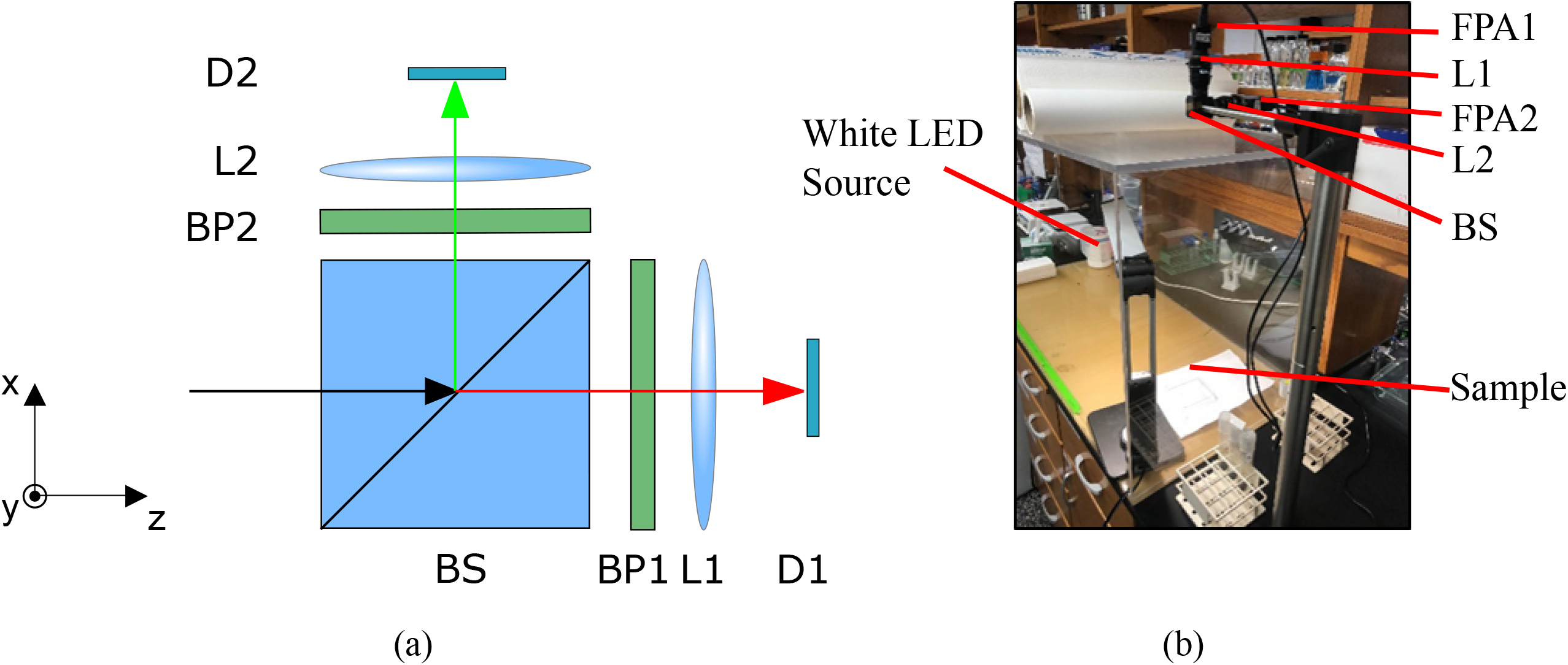
Band-pass filtered dual-camera system (a) schematic and (b) photograph.

In post-processing, an affine image registration algorithm [16] aligns the two images, taking into account translational and rotational differences. In addition, both of these images are normalized to flat field images taken prior to measurements with the same camera settings. This corrects for non-uniform spectral radiance from the LED and differences in shutter time and aperture size in the cameras. Finally, the calculation from Eq. (2.3) is applied using the spatially registered images as input arguments. To enable rapid segmentation of the individual leaf’s boundaries while maintaining moisture, they were placed on a red terrycloth background. This provided a high positive value of *v* that could be easily distinguished from the lower values produced by the diseased or healthy tissues. The background pixel values are set to global minimum values during post-processing so that the leaves in the contrast images stand out better visually.

## 3. Results

The multispectral camera system was used to obtain contrast images of two distinct species - *Lactuca sativa* cv. Salinas and *L. serriola* acc. US96UC23 (US96), parents of a recombinant inbred mapping population [17]. The leaves of each species were inoculated with *B. cinerea* and images acquired over several days as the symptoms spread. Figure 6 shows the contrast images of a US96 and Salinas leaf over seven days alongside a reference color image taken using a color USB camera. It can be seen that the healthy lettuce areas have lower values of *v* as compared to damaged areas, as expected from our hyperspectral results given healthy tissue yields a lower reflectance at 670 nm compared to 540 nm (producing a negative value), while infected tissue causes the reflectivity at both wavelengths to become more similar (producing a value closer to 0).

**Figure 6.**
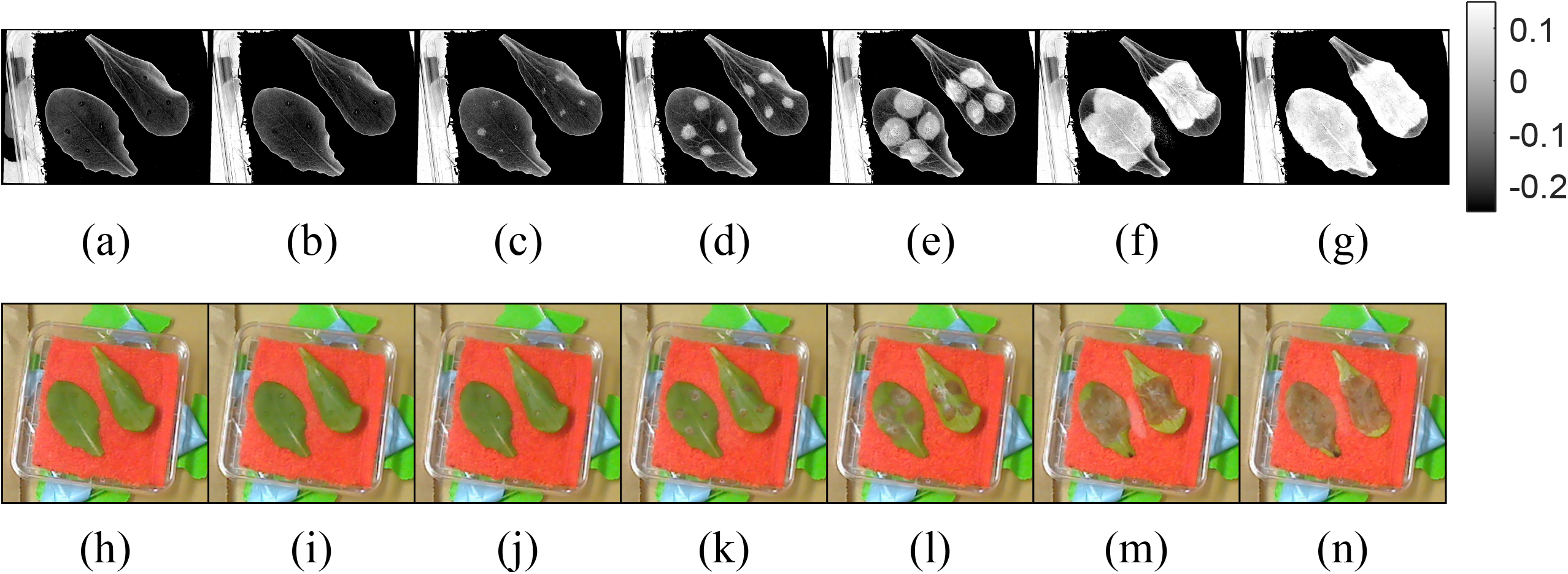
Contrast images (*v*) and color reference images taken over six days at roughly the same time each day of Salinas (left in images) and US96 (right in images) lettuce varieties inoculated with *Botrytis cinerea*. Each contrast image corresponds to the number of days after inoculation: (a) Day 0, (b) Day 1, Day 2, (d) Day 3, (e) Day 4, (f) Day 5, and (g) Day 6. Each color reference image also corresponds to a day after inoculation: (h) Day 0, (i) Day 1, (j), Day 2, (k) Day 3, (l) Day 4, (m) Day 5, and (n) Day 6.

Additional processing was implemented to measure the percent area of the lettuce leaves that were covered by the gray mold disease. Each leaf was segmented from the background by thresholding pixel values. By using a bright red background in the images, the contrast between the band-pass images was much higher than any measurements on a particular leaf. Using this relation, the leaves can be isolated from the image by classifying pixel values less than a specified background threshold as belonging to leaves. This was used to create a binary mask of the original contrast image that was used to isolate single leaves for further processing. Within each leaf, the diseased and healthy areas were segmented by further thresholding the pixel intensities. In order to determine a reasonable threshold to best discriminate between healthy and symptomatic lettuce areas, ROIs were labeled in a set of contrast images to compare the healthy and symptomatic values.

A histogram of *v* for both the healthy and symptomatic pixel classes are presented in Figure 7. Using the symptomatic and healthy means (*μ*_*s*_ and *μ*_*h*_) and standard deviations (*σ*_*s*_ and *σ*_*h*_), a decision threshold, *η*, was defined by estimating a probability density function (PDF) for each type and calculating the location where the two PDFs were equal. Based on this calculation, it was determined that a contrast threshold of *η* = −0.0915 was sufficient for discriminating healthy and symptomatic tissues.

**Figure 7.**
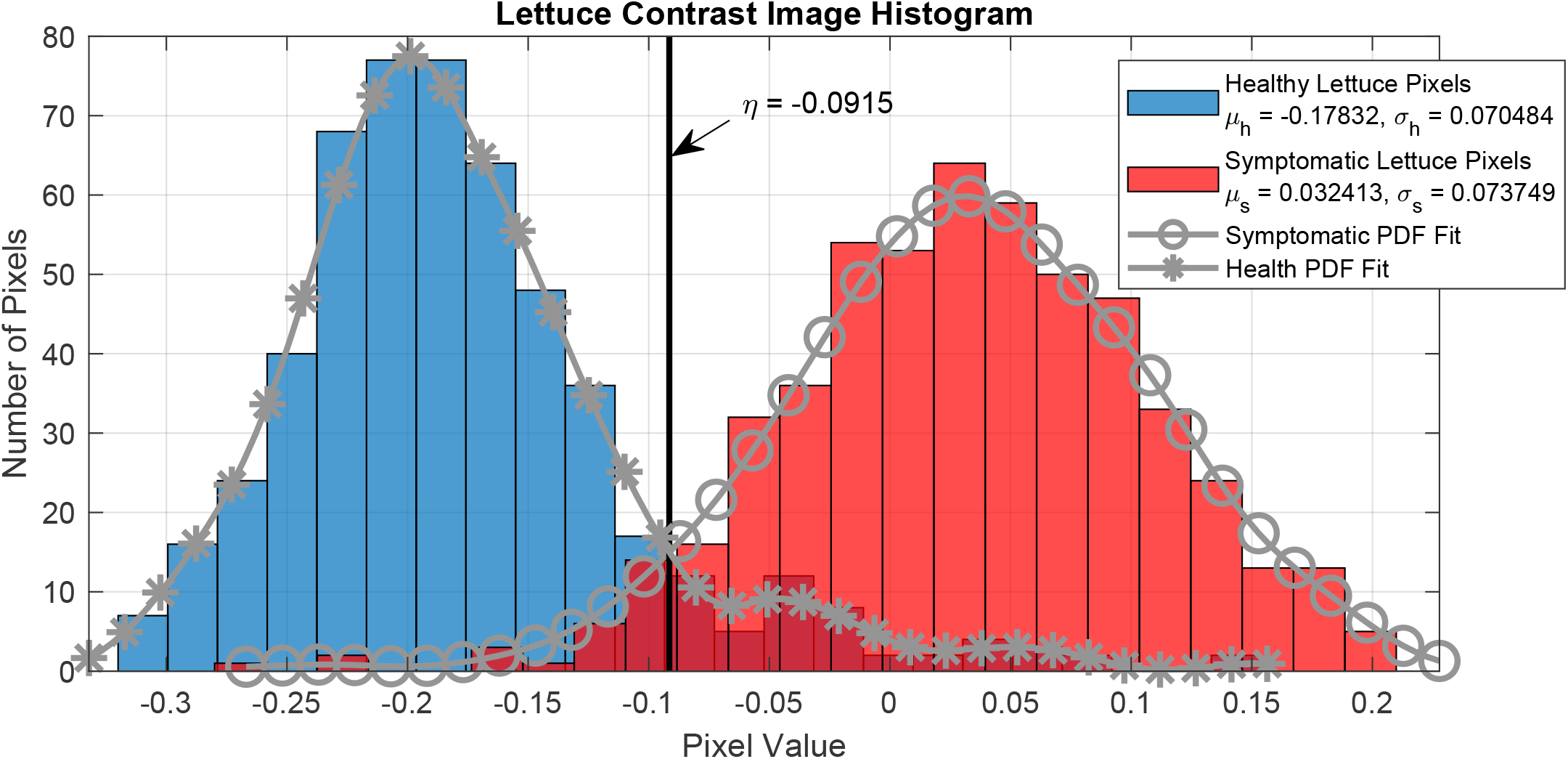
Histogram of pixel values corresponding to healthy and symptomatic lettuce areas compiled for both US96 and Salinas varieties. A threshold for discriminating between the two classifications was determined using these distributions.

In Figure 8, a receiver operating characteristic (ROC) curve was created based on the aforementioned PDFs. From this curve, it can be seen that for a probability of false positive classification, *P*_*F*_, of symptomatic tissue equal to 0.09316, the true positive probability, *P*_*D*_, is 0.9525.

**Figure 8.**
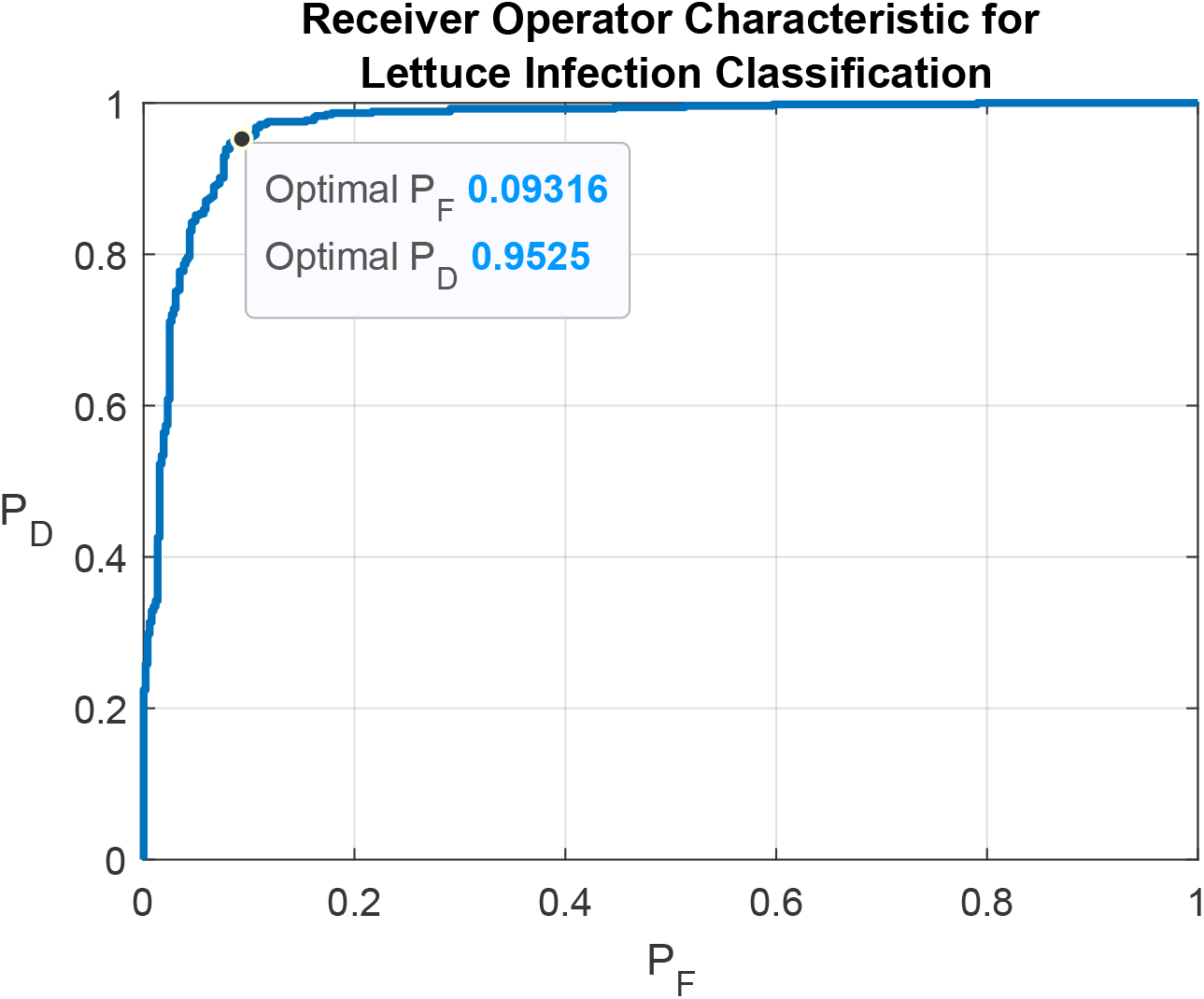
Receiver Operating Characteristic (ROC) for classifier based on thresholding band-pass contrast images of infected lettuce leaves. P_F_ is the false positive disease detection probability, while P_D_ is the true positive disease detection probability. The area under the curve is 0.9698.

A binary classification image that uses this decision threshold is depicted in Figure 9. There are some false-positive areas in the binary imagery that are indicated as belonging to a diseased lettuce region when they are not. These areas occur around the leaves’ edges where the red background was classified as part of the leaf due to shadowing effects and was not removed from the image, and along the leaf veins which can tend to have more yellow coloration than the leaf blades. To account for this when calculating the amount of disease symptoms on a particular leaf, an offset subtraction described by

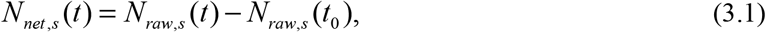

where *N*_*raw,s*_(*t*) is the number of pixels classified as symptomatic at time, *t*, and *t*_*0*_ is the time of the first binary image immediately of inoculation. *N*_*net,s*_(*t*) is the net number of symptomatic pixels in the binary image after *t* hours.

**Figure 9.**
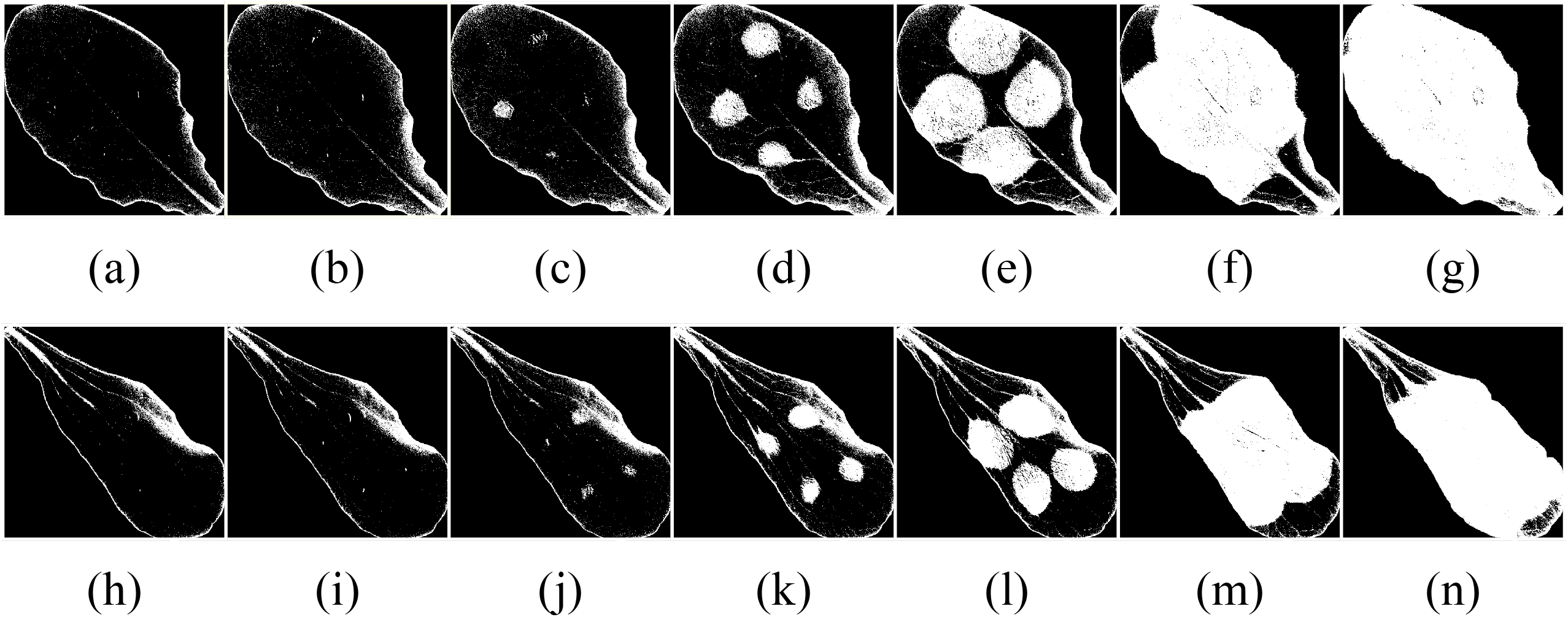
Timelapse binary images of Salinas and US96 leaves where white areas indicate disease spread detected while black areas indicate no symptoms detected. The classification decision was made by comparing the contrast images from the band-pass dual-camera system to a single threshold, *η*. Images are sequenced by days after *B. cinerea* inoculation for Salinas leaves on (a) Day 0, (b) Day 1, (c) Day 2, Day 3, (e) Day 4, (f) Day 5, and (g) Day 6, and for US96 leaves on (h) Day 0, (i) Day 1, (j) Day 2, (k) Day 3, (l) Day 4, (m) Day 5, and (n) Day 6.

The ratio of the diseased leaf area to the total leaf area was calculated and is presented in Figure 10 (a) and (b) for Salinas and US96, respectively. Both the diseased area percentage with and without offset subtraction in Eq. (3.1) are provided in the plots. This provides a data metric for studying the infection’s spread and leaf damage over time. According to these plots, the symptoms spread at what appears to be an exponential rate until the leaf is close to fully damaged. At this point, the rate of disease spread starts to decrease, consistent with a population growth curve approaching the carrying capacity.

**Figure 10.**
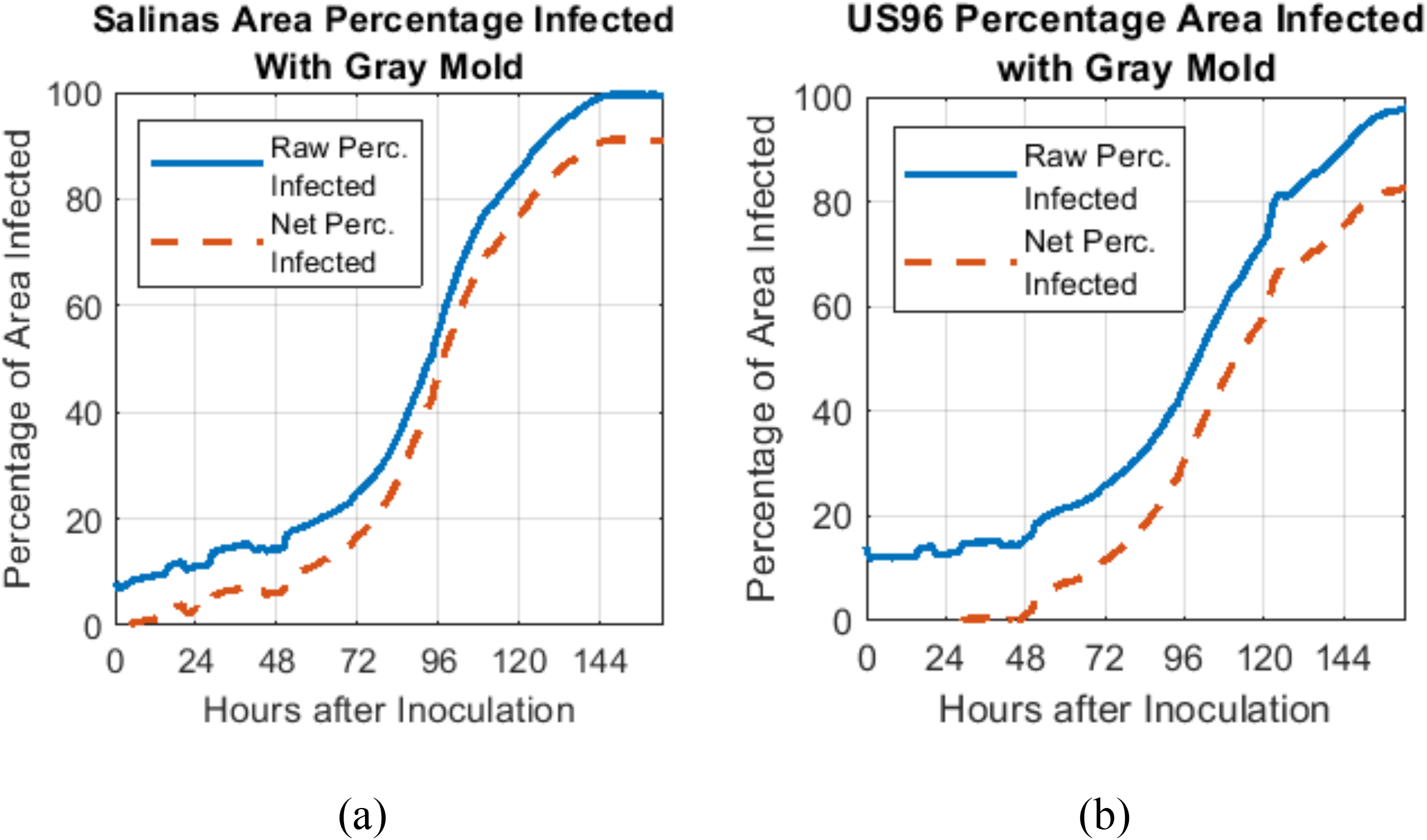
Plot of percentage leaf area infected with gray mold for (a) Salinas and (b) US96 types

## 4. Conclusion

A methodology for building a multispectral imaging system with the purpose of locating gray mold infection in plant leaves was demonstrated. A pushbroom hyperspectral imaging system was leveraged to select two narrow spectral band-pass filters to use in the dual-camera system. Post-processing of the collected data provides a basis for visually identifying infection areas and quantifying the spread over time. By taking the normalized difference between the two band-pass images, a significant contrast between leaf areas with gray mold disease and unaffected areas was observed, and the computation required little complexity. A simple threshold was used to programmatically discriminate between infected and healthy leaf areas with reasonable accuracy.

## 5. Acknowledgements

The authors would like to thank the United States Department of Agriculture (USDA) for funding this study as part of the National Needs Graduate Fellowships Program (2016-38420-25324: “Multidisciplinary Graduate Training in Advanced Technologies for High Yield Sustainable Agriculture”). We also appreciate the assistance from Katherine Denby, University of York, in helping establish *B.cinerea* infections, and Richard Michelmore, UC Davis, for providing the lettuce genotypes for the study.

## 6. Author Contributions

M.W.K. and C.J.D. conceived the original research experiment. C.G.S. wrote the original manuscript with edits and revisions provided by M.W.K., C.J.D., S.M.R, and C.G.S. S.M.R. and C.J.D. grew the lettuce plants, generated the *B. cinerea* suspension, and established and implemented the procedure for inoculating the lettuce leaves with the suspension for the experiments. C.G.S. performed the calibration for, and collected and processed data from the hyperspectral pushbroom camera with guidance from M.W.K. C.G.S. constructed the dual-camera band-pass system and processed and analyzed the data acquired from the system with guidance from M.W.K.

## Notes

### Competing Interest Statement

The authors have declared no competing interest.

